# The Photoconvertible Fluorescent Probe, CaMPARI, Labels Active Neurons in Freely-Moving Intact Adult Fruit Flies

**DOI:** 10.1101/2020.02.18.954669

**Authors:** Katie A. Edwards, Michael B. Hoppa, Giovanni Bosco

## Abstract

Linking neural circuitry to behavior by mapping active neurons *in vivo* is a challenge. Both genetically encoded calcium indicators (GECIs) and intermediate early genes (IEGs) have been used to pinpoint active neurons during a stimulus or a behavior but have drawbacks such as limiting the movement of the organism, requiring *a priori* knowledge of the active region or having poor temporal resolution. CaMPARI (calcium-modulated photoactivatable ratiometric integrator) was engineered to overcome these spatial-temporal challenges. CaMPARI is a photoconvertible protein that only converts from green to red fluorescence in the presence of high calcium concentration and violet (405 nm) light. This allows the experimenter to precisely mark active neurons within defined temporal windows. The photoconversion can then be quantified by taking the ratio of the red fluorescence to the green. CaMPARI promises the ability to trace active neurons during specific stimulus; however, CaMPARI’s uses in adult *Drosophila* have been limited to photoconversion during fly immobilization. Here, we demonstrate a method that allows photoconversion of multiple freely-moving intact adult flies during a stimulus. Flies were placed in a dish with filter paper wet with acetic acid (pH=2) or neutralized acetic acid (pH=7) and exposed to photoconvertible light (60 mW) for 30 min (500 ms on, 200 ms off). Immediately following photoconversion, whole flies were fixed and imaged by confocal microscopy. The red:green ratio was quantified for the DC4 glomerulus, a bundle of neurons expressing *Ir64a*, an ionotropic receptor that senses acids in the *Drosophila* antennal lobe. Flies exposed to acetic acid showed 1.3-fold greater photoconversion than flies exposed to neutralized acetic acid. This finding was recapitulated using a more physiological stimulus of apple cider vinegar. These results indicate that CaMPARI can be used to label neurons in intact, freely-moving adult flies and will be useful for identifying the circuitry underlying complex behaviors.

## 1 Introduction

Calcium is an essential secondary messenger in neurons and has proven to be one of the most widely used signals to trace neural activity (Palmer and Tsien, 2006). Most neurons have a low intracellular calcium concentration of 50-100 nM that rises 10-100 fold higher during neural activation (Berridge et al., 2000), thus activity can be reliably marked by a spike in calcium concentration. Initially, measuring intracellular calcium was dependent on the delivery of dyes or fluorescent calcium indicators. While many of these indicators have good signal-to-noise ratios and fast kinetics (Paredes et al., 2008), they are typically only useful for a short time period after delivery (Jia et al., 2011). Protein-based genetically encoded calcium indicators (GECIs) were developed to allow for multiple readings over multiple days (Looger and Griesbeck, 2012). Over the past two decades, GECIs have been continuously developed and modified to improve brightness (Tian et al., 2009; Chen et al., 2013; Barnett et al., 2017), kinetics (Chen et al., 2013; Sun et al., 2013), stability (Tian et al., 2009; Chen et al., 2013) and sensitivity (Chen et al., 2013; Barnett et al., 2017) thus enabling more efficient tracking of changes in activity over time in neuronal subpopulations and whole brains in a variety of model organisms.

Despite the broad success of GECIs in in vivo imaging, tracking neural activity with behavior is limited due to the necessity to head-fix organisms to obtain high resolution images (Tian et al., 2009; Sun et al., 2013) with the exception of the development of the miniscope for rodents and other small mammals (Aharoni and Hoogland, 2019). However, the field of view is often limited by the necessity to deliver light into the brain and may require *a priori* knowledge of the specific brain region relevant to the specific stimulus, regardless of whether the animal is head-fixed under a microscope or freely-moving with a miniscope (Aharoni and Hoogland, 2019; de Groot et al., 2020). Because some circuit activity is so short-lived, it may escape detection unless the experimenter has specific knowledge of where and when to image. As such, GECIs have limited use as a circuit-discovery tool in identifying activated circuits in freely behaving animals. This is especially true in non-transparent organisms or systems with non-transparent developmental stages. A potential solution to these challenges would be the ability to accurately “tag” or identify neurons during a defined stimulation period allowing detailed imaging after the activity. To do just that, Fosque and colleagues invented a calcium integrator, which combined a photoswitchable protein (mEOS) with molecular aspects of GCaMP, a GECI, to make photoswitching dependent on both calcium and violet light (Fosque et al., 2015). With this new tool, active neurons expressing this integrator would convert fluorescent emission from green to red only in the simultaneous presence of high calcium concentration from neural activity and user-supplied violet light. This calcium integrator was named CaMPARI (calcium-modulated photoactivatable ratiometric integrator). Because of the transient nature of calcium and the user defined photoconverting light, CaMPARI has tight temporal window to take snapshots of neural activity during defined behavior.

Thus far CaMPARI has been successfully deployed in behaving solitary juvenile zebrafish (Moeyaert et al., 2018; Sha et al., 2019) and fruit fly larva (Patel and Cox, 2017) due to the translucent nature of those organisms. However use in adult fruit flies has been limited due to the head cuticle of the fly. Most published reports of CaMPARI’s uses in the adult fruit fly, require the fly to be fixed or tethered on a microscope with direct illumination from a laser (Bohra et al., 2018; Manjila et al., 2019). While such experiments are valuable, they limit the movement of the fly and again require the animal to be solitary, which can affect how an animal responds to stimulus naturally (Ramdya et al., 2017). Thus, this is a central limitation of CaMPARI’s versatility to identify circuits involved in natural behaviors and in large scale experiments on numbers of flies used in each experiment, precluding a tremendous benefit of the *Drosophila* system.

Here we propose a setup that utilizes inexpensive and commonly used materials to take advantage of existing CaMPARI fly stocks in order to label neuronal activation in freely moving individual or groups of flies. This involves the use of standard Petri dishes with passive humidification and a series of violet LEDs that allow for photoconversion of CaMPARI in the freely-moving intact adult fly. With this setup, we demonstrate that photoconversion of specific neuronal circuits is possible in multiple sets of fruit flies simultaneously. We provide quantified photoconversion of an olfactory glomeruli known to be activated by exposure to acids (Ai et al., 2010) as a proof of principle that this method allows for quantifiable significant photoconversion through the intact head cuticle of the freely-moving adult fly. Because this method is scalable to any group size, we suggest that CaMPARI in *Drosophila* is easily adaptable for large scale genetic screens and/or discovery and mapping of novel circuits that are activated by the investigator’s stimulus of choice.

This method has the benefits of photoconverting multiple flies at once, allowing for the full movement and exploration behaviors of the fly, including social and group behaviors. We suggest that this approach will be useful in studying ecologically relevant behaviors, neurogenetic factors and the circuits that govern such behaviors.

## 2 Materials and Equipment

### 2.1 Fly Stocks

**UAS-CaMPARI-GAL4.** w[*]; P{y[+mDint2] w[+mC]=UAS-CaMPARI}attP40; PBac{y[+mDint2] w[+mC]=GMR57C10-GAL4}VK00040 PBac{GMR57C10-GAL4}VK00020 (Bloomington *Drosophila* Stock Center, catalog number: 58763).

**UAS-CaMPARI.** w[*]; P{y[+t7.7] w[+mC]=UAS-CaMPARI}attP40 (Bloomington *Drosophila* Stock Center, catalog number 58761).

**Ir64a-GAL4.** w[*]; P{w[+mC]=Ir64a-GAL4.A}183.8; TM2/TM6B, Tb[1] (Bloomington *Drosophila* Stock Center, catalog number 41732).

**Ir64a RNAi.** w1118; P{GD2314}v9011 (Vienna *Drosophila* Resource Center, catalog number 9011).

### 2.2 Fly Crosses

**UAS-CaMPARI x Ir64a-GAL4.** 10 virgin female w[*]; P{y[+t7.7] w[+mC]=UAS-CaMPARI}attP40 crossed with 5 male w[*]; P{w[+mC]=Ir64a-GAL4.A}183.8; TM2/TM6B, Tb[1]

**UAS-CaMPARI-GAL4 x Ir64a-RNAi** (Ir64a knockdown in nSyb neurons). 10 virgin female w[*]; P{y[+mDint2] w[+mC]=UAS-CaMPARI}attP40; PBac{y[+mDint2] w[+mC]=GMR57C10-GAL4}VK00040 PBac{GMR57C10-GAL4}VK00020 crossed with 5 male w1118; P{GD2314}v9011

### 2.3 Solutions

**5% (v/v) Acetic acid.** 5mL Glacial Acetic Acid (Fisher Chemical Acetic Acid, Glacial Lot# 185624) diluted to 100 mL with H_2_O. For neutralized acetic acid, neutralized to pH=7 with Sodium Hydroxide pellets (Fisher Scientific Lot# 115786).

**Apple Cider Vinegar** Bought from a local supermarket. For neutralized apple cider vinegar, neutralized to pH=7 with Sodium Hydroxide pellets (Fisher Scientific Lot# 115786).

**Phosphate-buffered saline (PBS).** 80g Sodium Chloride (NaCl) (EMD Chemicals SX0420-1 Lot# UG25BZEMS), 2g Potassium Chloride (KCl) (Fisher Scientific P217-500 Lot# 117211), 14.4g Sodium Phosphate, Dibasic Anhydrous (Na_2_HPO_4_) (Fisher Chemical S374-500 Lot# 151771), 2.4g Potassium Phosphate, Monobasic (KH_2_PO_4_) (EMD Chemicals PX1565-1 Lot# 2011061522) diluted to 1L with H_2_O, then sterilized by autoclave to make a 10x solution. 1X PBS is made by diluting 100 mL 10X PBS to 1L with H_2_O.

**PBT (PBS + 0.1% Triton-X 100).** 100 mL 10X PBS and 10 mL Triton-X 100 (Sigma Aldrich Lot# MKBW8386V) diluted to 1L with H_2_O.

**4% formaldehyde in PBT.** 2 mL 16% formalehyde solution (w/v), Methanol-free (Thermo Scientific Lot# UC2742481) and 6 mL PBT.

**Vectashield Antifade Mounting Medium** (Vector Vectashield Lot# ZC0829)

### 2.4 Photoconversion Equipment

Sanworks Pulse Pal v2 (Sanders and Kepecs, 2014) (can be substituted with any pulse generator)

405 nm LEDs (Thorlabs M405L3)

Collimation Collars (Thorlabs SM1P25-A)

Current Controller (Thorlabs T-cube LED Driver LEDD1B)

Adaptor Plate for current controllers (KAP101)

Power Supply for current controllers (Thorlabs KCH601)

Analog Handheld Power Meter Consol (Thorlabs PM100A)

Standard Photodiode Sensor for 400 – 1100nm (SC121C)

### 2.5 Software

Zeiss Zen Black 2.3

Fiji (Schindelin et al., 2012)

Graphpad Prism 8.3.0

### 2.6 Other

Petri dishes (35 × 10 mm) (Falcon 351007)

42 Ashless Filter Paper (70 mm diameter) (GE Healthcare Whatman 1442-070 Lot# 16946862)

“PTFE” Printed 10-well 6 mm well diameter Slides (Electron Microscopy Sciences 63434-06 Lot# 171020)

20×50 mm No. 1 1/2 Coverslips (VWR 48393 194)

## 3 Procedures

### (A) *Drosophila* cultures, animal manipulation and genetic crosses

**Note:** *Drosophila melanogaster* was used for all experiments described and culture conditions were used as described below. One notable exception to standard fly manipulation was the use of ice to anesthetize adults, instead of the usual humidified carbon dioxide. This was especially important for adult flies that were subsequently used in photoconversion experiments. Exposure to CO_2_ could activate olfactory as well as acid sensing neurons while also possibly triggering a stress response (Suh et al., 2004; Ai et al., 2010).

1. Drosophila cultures were reared on standard cornmeal-molasses media in plastic bottles (Genesse, catalog number 32-130) at 22°C, ∼40% humidity and 12hr-dark/12hr-light cycle.
2. Genetic crosses were necessary in order to achieve some genotypes of interest, for example where CaMPARI was being expressed in specific regions of the brain or when short-hairpin RNA (shRNA) was being expressed in addition to CaMPARI. Typically, 10 virgin females 4-6 days post-ecolsion (dpe) were crossed with 5 males and placed in a 22mm food vial (Genesee, catalog number 32-116). Adult parents were discarded after 4-7 days or transferred into a fresh food vial in order to continue laying eggs. Controlling the number of females and the number of days the females are allowed lay eggs ensures that progeny are not overcrowded, thus ensuring maximum consistency in access to nutrition during development and adult body and tissue size. Adult progeny were collected for photoconversion 14-18 days after the initial cross was started.
  - Virgin UAS-CaMPARI flies were crossed to Ir64a-GAL4 flies at 4 dpe and progeny collected at 1-2 dpe and isolated in groups of 5 females and 3 males at 25°C.
  - Pan-neuronal expressing CaMPARI flies were collected at 1-2 dpe and isolated in groups of 5 females and 3 males at 25°C. Since this genotype was maintained as a stable stock it was not necessary to perform a genetic cross.
  - In order to produce adult flies expressing CaMPARI and Ir64a-RNAi in nSyb-expressing neurons 10 virgin females were crossed to 5 UAS-Ir64a-RNAi males and progeny collected at 1-2 dpe and isolated in groups of 5 females and 3 males at 25°C.

### (B) Photocoversion setup (duration: 1 d)

The setup can be adjusted to suit the needs of the investigator/experiment. In **Figure 1**, we show our setup of 16 LEDs (Thorlabs M405L3) with collimation collars (Thorlabs SM1P25-A) adjusted to match the area of the Petri dishes. Each LED is connected to a current controller (Thorlabs T-Cube LED Driver LEDD1B) where the power of the LED can be adjusted via the current while the power can be measured for each LED using an optical power meter (Thorlabs PM100A) and a standard photodiode sensor (SC121C). The current controllers can be powered individually but are here powered by a power supply (KCH601) with an adaptor plate (KAP101). Each current controller is connected to a pulse generator (Sanworks Pulse Pal v2) with standard female BNC to female BNC cables.

**Figure 1:**
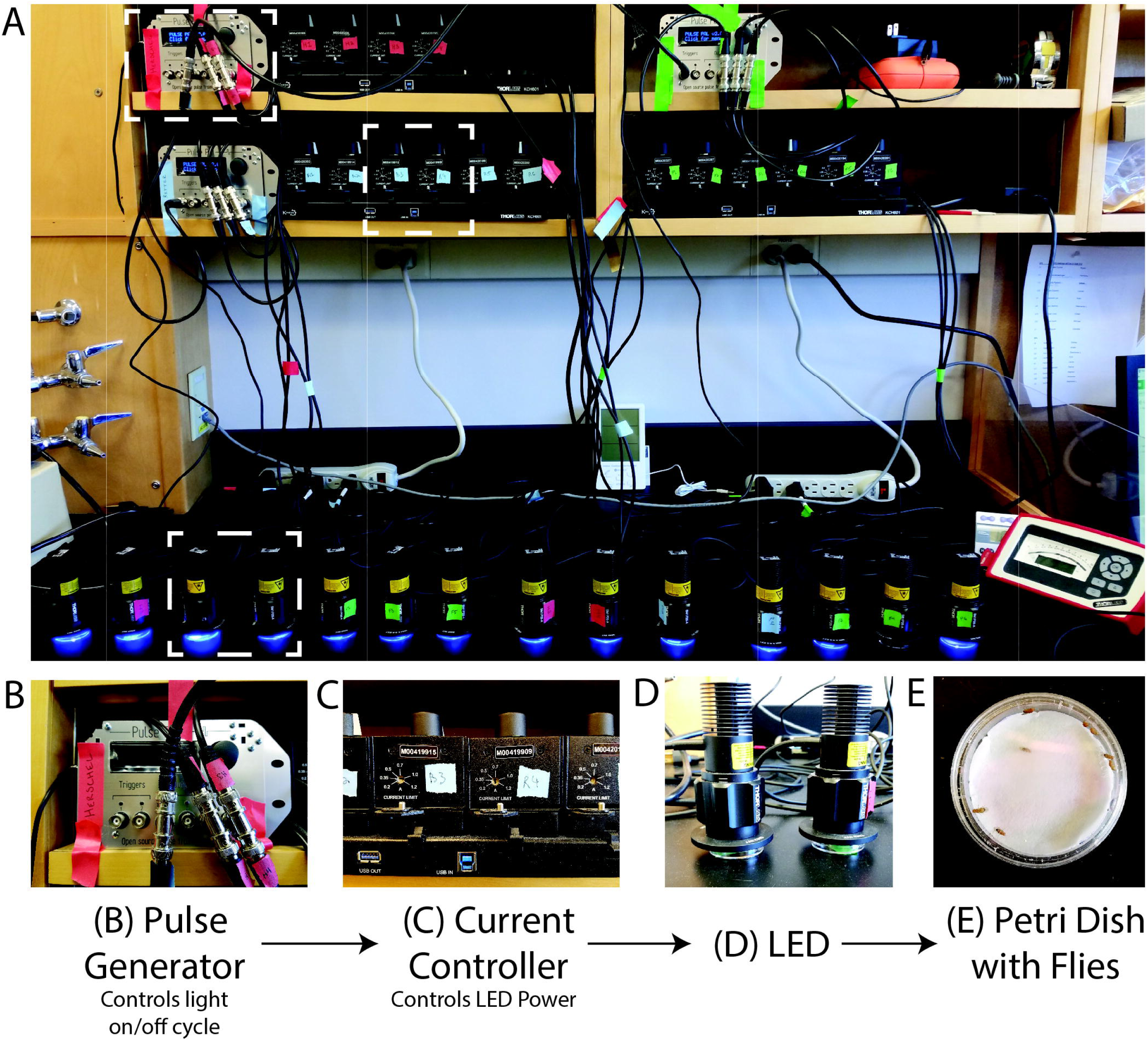
Photoconversion Setup. A view of the setup as used to induce photoconversion in freely moving flies. **(A)** Sanworks Pulse Pal v2 is used to control the LED on/off cycle. **(B)** The pulse generators are connected to current controllers (Thor Labs M00420201). The knobs of the current controllers are used to set the LED power. **(C)** The 405 nm LEDs (Thor Labs M405L3) light is dispersed with collimation collars (Thor Labs SM1P25-A) set to match the area of the dishes. **(D)** Petri dishes are prepared with moistened filter paper and can contain individual or sets of adult *Drosophila* that sit underneath the LEDs.

The PulsePal can be programmed to control the on/off cycle of the LEDs. The PulsePal has four “output channels” that can be programmed separately and triggered simultaneously by linking them to the same trigger channel.

### (C) Photoconversion of flies (duration: 1 h)

1. Prepare the Petri dishes that will contain the flies during photoconversion. Trim ashless filter paper by trimming to the size of the Petri dish. Moisten the filter paper by pipetting directly onto the paper 150 µL PBS or other solutions to be used as stimuli. Drops should be evenly distributed so that the entire filter paper is moistened. The paper should be pushed flat to the bottom of the dish so that flies cannot get into crevices underneath the paper. PBS can be substituted with solutions to be used as stimuli. Here, as experimental conditions we used 5% acetic acid and apple cider vinegar solutions in place of PBS. **Critical step:** It is necessary to introduce a solution into the dish to control the humidity within the dish. Without added solution, the flies can overheat and desiccate, killing them during photoconversion.
2. Place vials containing flies on ice for 20 minutes. Vials are partially submerged in a bucket containing ice. **Note:** Certain anesthetizing steps may induce neural activation, as described above.
3. Calibrate LED power by adjusting the dial of the current controller using a power meter. In this study, we used 75 mW/cm^2^ ± 0.5 mW/cm^2^ power. **Note:** LED power may need to be adjusted for deeper brain regions or neurons with less frequent firing rates.
4. Setup pulse generator (**Figure 1**). The Pulse Pal was programmed to deliver monophasic pulses of 5 V every 500 ms with a phase interval of 200 ms on a continuous cycle until manually aborted. **Note:** In our setup, we used an open source hardware, the Sanworks Pulse Pal, however any pulse generator can be used that is compatible with the LEDs.
5. From the anesthetized fly vials that were placed on ice, dispense 5 female and 3 male adult (4-6 dpe) into each dish prepared in Step 1 and place lids on each dish. **Critical step:** Wait until flies are actively moving on their own before moving to the next step. This ensures they have fully recovered from anesthesia.
6. Place LEDs on top of dishes and set PulsePal/pulse generator to “Continuous”. Allow the program to continue for 30 minutes or desired time.
7. Prepare fixation reagents (4% formaldehyde in PBT) and dispense in 800 µL aliquots in 1.5 microcentrifuge tubes.
8. End photoconversion protocol. Anesthetize the flies by delivering CO_2_ into the space between the dish and the lid using a syringe. Transfer the whole flies directly into the 1.5 mL tubes containing fixation solution. Spin down the tubes containing flies in a centrifuge for 5 min at 13,000 rpm. Store flies in fixation solution at 4°C for 14-18 hrs. **Note:** Alternatively, fixation is not necessary as the tissue could be imaged live or dissected and then fixed. CaMPARI is sensitive to fixation and so signal, particularly red signal, is compromised (Fosque et al., 2015). Keep in mind turnover rate of photoconverted CaMPARI, which is ∼5 hrs, when performing live imaging or dissections. Recently an antibody has been developed that specifically binds the red CaMPARI that can be used to amplify the signal (Moeyaert et al., 2018).

### (D) Brain dissection and tissue sample preparation (duration: 2 h)

1. Dissect brain from head cuticle, as previously described (Wu and Luo, 2006) into the wells of a glass spot plate.
2. Wash dissected brains three times for 10 min by pipetting 150 µL PBT into each well containing tissue.
3. Wash dissected brains four times for 5 min by pipetting 150 µL PBS into each well containing tissue.
4. Dispense Vectashield Antifade Mounting Medium into each well of a slide enough to just fill the well.
5. Carefully transfer dissected brains into the wells of the slide. Each well should contain no more than three brains. **Critical step:** Brains should be oriented such that the region of interest is closest to the coverslip to prevent light dispersal through tissue and obtain better resolution images.
6. Mount brain tissue on slides with Vectashield Antifade Mounting Medium. **Note:** This mounting medium remains a liquid on the slide and it’s recommended to seal the slide.

### (E) Imaging (duration: 2-8 h)

Images were collected using a Zeiss LSM 880 microscope and Zeiss Zen Black 2.3 software using a 20x Plan-apochromatic (0.8 NA) objective. 488 nm and 561 nm laser lines were used for illumination and a GaAsP detector for imaging. 1024×1024 images were collected with a 1.02 µs pixel dwell and 16 bit depth. Z-stacks were collected at a 0.97 µm step.

### (F) Image analysis (duration: 2-6 h)

1. For each sample, identify slices that contain the region(s) of interest (ROIs) using Fiji.
2. Combine slices and collapse into a single image using an average intensity projection (AIP) (**Figure 2C**).

**Figure 2:**
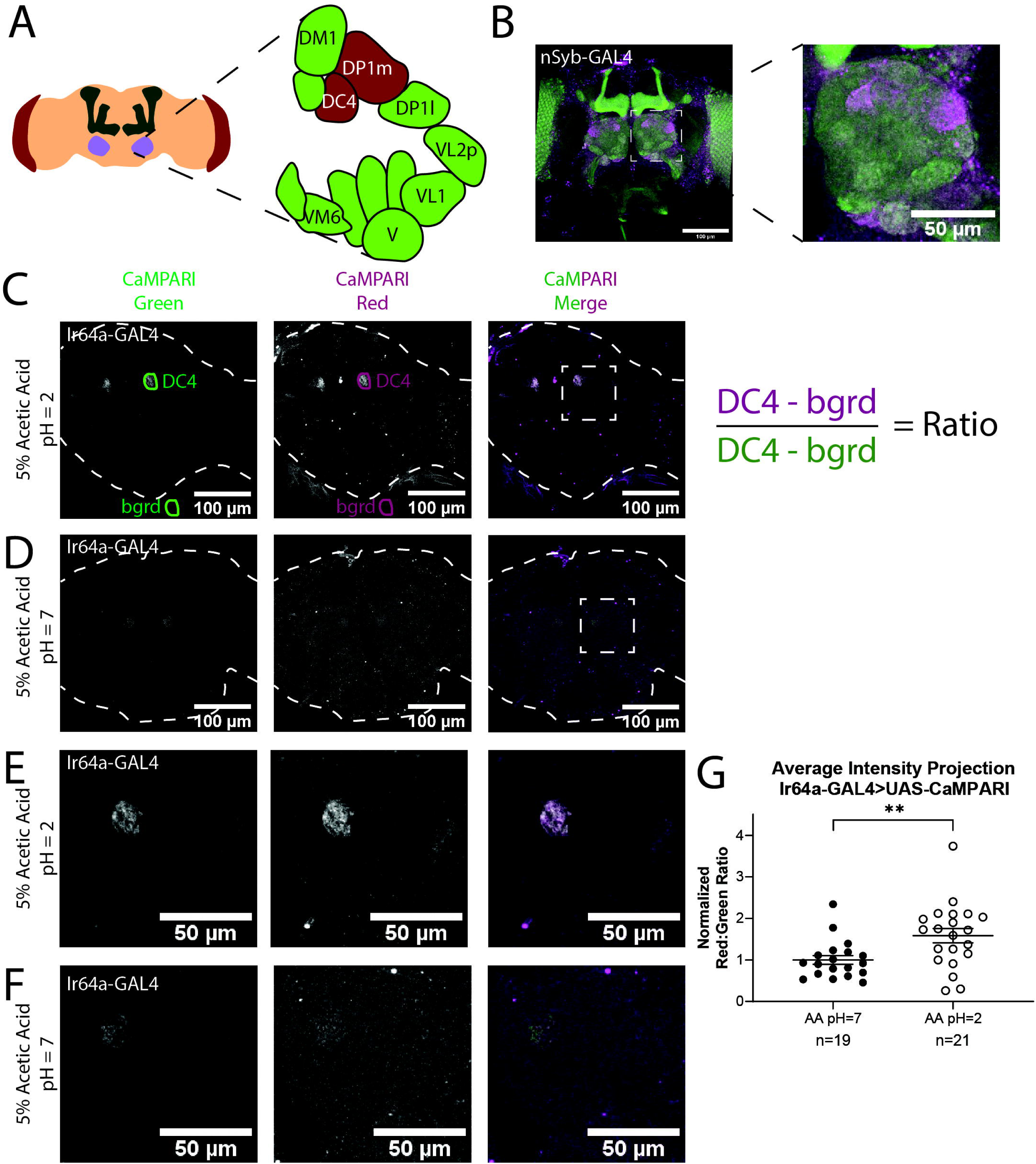
Analysis of Images of Flies Expressing CaMPARI. **(A)** Schematic of the *Drosophila* head labeling the antennal lobe and antennal lobe glomeruli in the region of interest. Acid-sensing glomeruli (DC4 and DP1m) are labelled in red. **(B)** Representative maximum intensity projection (MIP) of adult fly brain expressing CaMPARI pan-neuronally exposed to 5% acetic acid at pH=2 for 30 min and imaged at 20x. A white dotted square indicates a single antennal lobe blown up to the right demonstrating photoconversion of DC4 glomerulus. **(C-D**) Representative average intensity projection (AIP) images of adult fly brains expressing CaMPARI in Ir64a-expressing neurons exposed to 5% acetic acid (AA) at pH=2 **(C)** or pH=7 **(D)** for 30 min and imaged at 20x. A ROI was drawn around the DC4 glomeruli in the green channel (DC4) and then copied and moved to the background (bgrd) in order to background-subtract the average green intensity in the DC4 glomerulus. The same pair of ROIs were copied to the red channel to collect the background-subtracted average red intensity. The ratio was then collected by dividing the red intensity by the green intensity as shown on the right. Photoconversion is seen in the red channel in **(B)** when pH=2 but not in **(C)** when pH=7. The antennal lobes are blown up in **E-F** as designated by the white dotted rectangle. **(E-F)** Representative maximum intensity projection (MIP) images of the adult fly antennal lobes expressing CaMPARI in Ir64a-expressing neurons exposed to 5% AA at pH=2 **(E)** or pH=7 **(F)** for 30 min. **(G)** The Red:Green ratio of flies exposed to AA of pH=2 and pH=7 for 30 minutes. Flies exposed to AA pH=2 (n=21) have 1.6-fold greater photoconversion compared to flies exposed to AA pH=7 (n=19). **P=0.0064, Mann-Whitney U-test
3. Use the “polygon tool” to trace the ROI within the AIP in the green channel, as in “DC4” in **Figure 2C**.
4. Copy the ROI and move to a region outside the imaged brain, as in “bgrd” in **Figure 2C**. This ROI will be used for background subtraction in the green channel.
5. Copy the same exact ROIs to the AIP of the red channel.
6. Use the “Measure” function to calculate the average intensity of each ROI.
7. Subtract background fluorescence from each ROI. For example, the average intensity of “bgrd” ROI is subtracted from the average intensity of the “DC4” ROI.
8. Divide background-subtracted red channel fluorescence by background-subtracted green channel fluorescence to determine the ratio.
9. Repeat steps 3-8 for any additional ROIs in the brain.
10. Red:green ratios were normalized to the ratios of images from flies exposed to pH=7 solutions by grouping flies that were photoconverted simultaneously. **Critical step:** It is important to only compare images that were taken on the same day to avoid batch effects. Microscope laser power can vary day-to-day and affect the quantification of the images.

### (G) Statistics

All data are presented as means ± SEM. Comparisons between two groups were performed using a Mann-Whitney U test in Graphpad Prism 8.3.0.

## 4 Results

### 4.1 Acetic acid exposure results in specific photoconversion of the DC4 glomerulus in flies expressing CaMPARI in Ir64a-expressing neurons

We first demonstrated that we could get specific photoconversion in acid-sensing neurons through the head cuticle of freely-moving adult flies by exposing flies with pan-neuronally expressing CaMPARI to 5% acetic acid (AA) (pH=2). Previously it was shown that Ir64a-expressing neurons of the DC4 glomerulus (**Figure 2A**) are specifically activated by acids (Ai et al., 2010). Adult intact freely-moving flies were exposed to 5% AA pH=2 for 30 minutes while also being exposed to photoconvertible light (405 nm; 500 ms on, 200 ms off). This resulted in a photoconverted region within each antennal lobe consistent with the DC4 glomerulus (**Figure 2B**).

To confirm this photoconverted region was specific to the acid-sensing neurons of the DC4 glomerulus, we limited expression of CaMPARI to the DC4 glomerulus by driving expression of CaMPARI with an Ir64a GAL4 driver (**Figure 2C-F)**. Intact, freely-moving adult flies were exposed to 5% AA (pH=2) or neutralized 5% AA (pH=7) for 30 minutes while simultaneously exposed to photoconvertible light. Average intensity projection (AIP) images of the full fly brain are shown for 5% AA (pH=2) (**Figure 2C**) and 5% AA (pH=7) (**Figure 2D**). As shown in the green channel, expression of CaMPARI is limited to the DC4 glomerulus. The signal of photocnverted CaMPARI is present for 5% AA (pH=2), but not when the acetic acid has been neutralized. Maximum intensity projection (MIP) of a single antennal lobe, as designated by the white dotted box in **Figure 2C** and **2D**, are shown for both 5% AA (pH=2) (**Figure 2E**) and neutralized AA (pH=7) (**Figure 2F**). These results confirm that photoconversion of CaMPARI in intact, freely-moving adult flies exposed to acetic acid is specific to the DC4 glomerulus.

### 4.2 Photoconversion of CaMPARI can be quantified in freely-moving intact adult flies

One of the major benefits to using CaMPARI as a tool for tracing active neurons is its use as a ratiometric integrator, meaning the level of photoconversion can be quantified by taking the ratio of CaMPARI intensity in the red channel to the CaMPARI intensity in the green channel. We quantified the red:green ratios of average intensity projection (AIP) images of freely-moving intact adult flies expressing CaMPARI in Ir64a-expressing neurons exposed to 5% acetic acid (AA) (pH=2) (**Figure 2C**) or neutralized 5% AA (pH=7) (**Figure 2D**). Five sets of 3-5 flies were exposed to acetic acid or neutralized acetic acid for 30 minutes while simultaneously exposed to photoconvertible light (405 nm; 500 ms on, 200 ms off). Immediately following photoconversion, flies were fixed in 4% formaldehyde in PBS + 0.1% TritonX-100 overnight. Brains were dissected from female flies and mounted on slides and imaged by laser-scanning confocal microscopy.

AIP images were constructed with Fiji using Z-project for all slices containing CaMPARI expression. The “polygon tool” was used to draw a region of interest (ROI) around each glomeruli in the green channel as shown in **Figure 2C-D** (“DC4”). These ROIs were saved to the ROI Manager in Fiji then each ROI was copied and moved to a region outside the fly brain to collect a background measurement (“bgrd”) for the green channel. All four of these ROIs were then copied to the red channel. The ratio was then calculated as the “DC4” intensity of the red channel minus red channel “bgrd” intensity divided by the “DC4” intensity of the green channel minus the green channel “bgrd” intensity. This resulted in two ratios for each fly brain, one for the right “DC4” glomeruli and a second for the left. Because these ratios were always comparable they were averaged to give a single datum point for each fly brain (**Figure S1**). Each ratio was then normalized to the average ratio of flies exposed to 5% AA (pH=7) for each set that was photoconverted and imaged on the same day to minimize batch effects. We found that flies with exposure to 5% AA (pH=2) (n=19 individual flies) had 1.6-fold photoconversion compared to flies exposed to 5% AA (pH=7) (n=21 individual flies) in the DC4 glomerulus (**Figure 2G**).

To confirm measurements taken from AIP images accurately reflect each slice of the ROI, ROIs were drawn for every slice and averaged together. These ratios were not appreciably different from the ones measured from AIP images (**Figure S2**), thus we continued to use only AIP images to quantify photoconversion.

These results confirm we can use AIP images to quantify the photoconversion ratio of a ROI in the fly brain when CaMPARI is photoconverted in the freely-moving intact adult fly.

### 4.3 Photoconversion of CaMPARI is comparably quantified in freely-moving intact flies expressing pan-neuronal CaMPARI

While it is useful to limit the expression of CaMPARI to subsets of neurons to obtain photoconversion ratios, for behaviors that recruit multiple regions of interest it may be more useful to quantify CaMPARI when it is expressed pan-neuronally. We wanted to confirm that ratios calculated from AIP images of pan-neuronally expressed CaMPARI would be comparable to the ratios we calculated when CaMPARI expression was limited to Ir64a-expressing neurons. Five sets of 3-5 flies expressing CaMPARI driven by a synaptobrevin (nSyb) GAL4 driver were exposed to 5% acetic acid or neutralized 5% acetic acid for 30 minutes while simultaneously exposed to photoconvertible light (405 nm; 500 ms on, 200 ms off) and their brains dissected and imaged as described above. Maximum intensity projection (MIP) images of one antennal lobe are shown in **Figure 3** for flies exposed to either 5% AA (pH=2) (**Figure 3A**) or neutralized 5% AA (pH=7) (**Figure 3B**). An arrow indicates the expected photoconversion in the DC4 glomerulus when the flies are exposed to 5% AA (pH=2). Ratios were calculated from AIP images as described above. We found freely-moving intact adult flies with pan-neuronally expressed CaMPARI exposed to 5% AA (pH=2) (n=20 individual flies) had 1.6-fold photoconversion compared to flies exposed to 5% AA (pH=7) (n=20 individual flies) in the DC4 glomerulus (**Figure 3C**), the same fold photoconversion as demonstrated in flies expressing CaMPARI in Ir64a-expressing neurons. This suggests that comparable photoconversion ratios can be calculated from either subsets of neurons expressing CaMPARI or pan-neuronally expressing CaMPARI.

**Figure 3:**
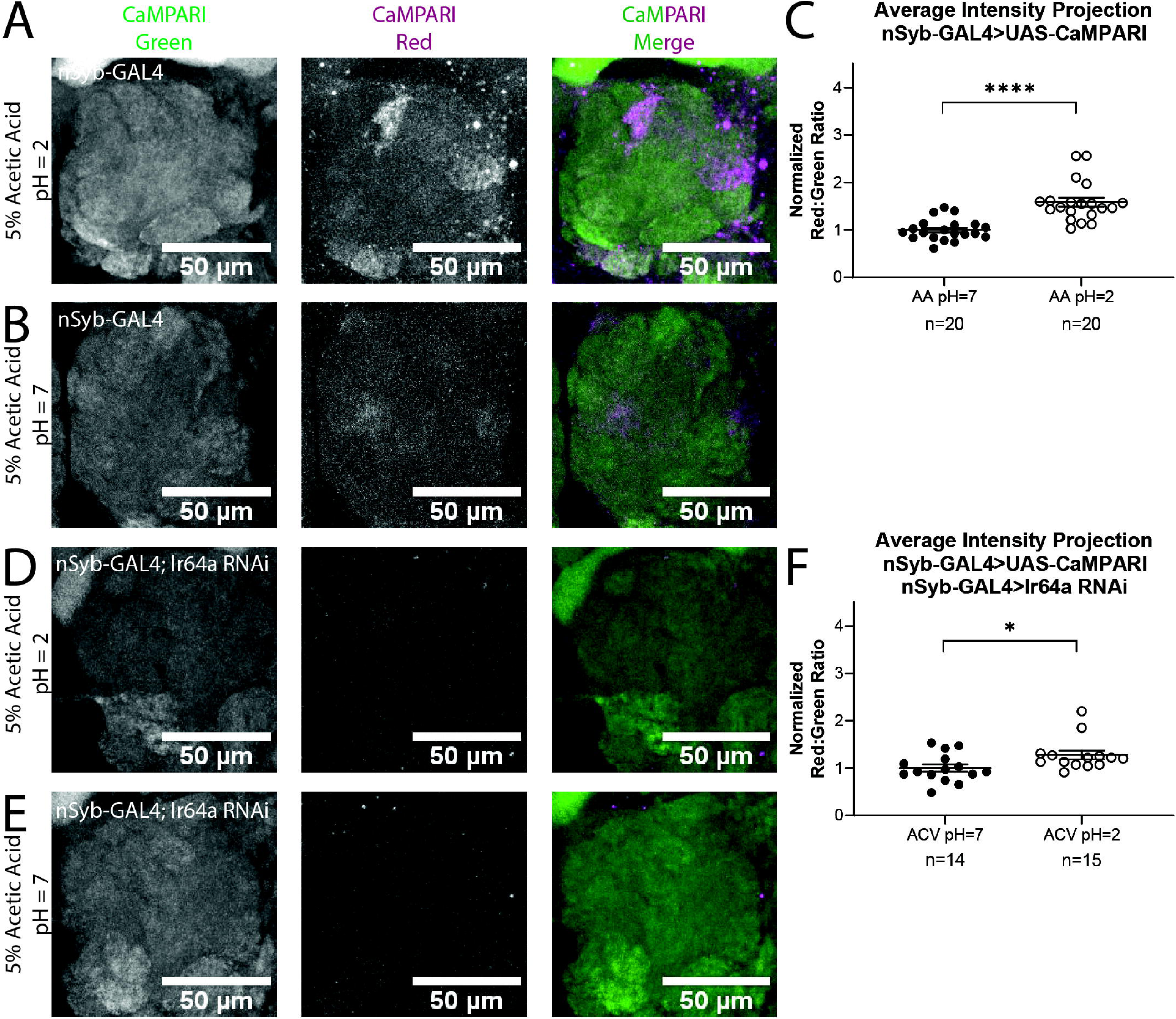
Flies with Pan-neuronally Expressed CaMPARI Have Comparable Measurable Photoconversion in Flies Exposed to Acetic Acid. **(A-B)** Representative MIP images of adult fly antennal lobe expressing CaMPARI in nSyb-expressing neurons exposed to 5% acetic acid (AA) at pH=2 **(A)** or pH=7 **(B)** for 30 min. Photoconversion is seen in the red channel in **(A)** when pH=2 but not in **(B)** when pH=7. **(C)** The Red:Green ratio of flies exposed to AA of pH=2 and pH=7 for 30 minutes. Flies exposed to AA pH=2 (n=20) have 1.6-fold greater photoconversion compared to flies exposed to AA pH=7 (n=20). ****P<0.0001, Mann-Whitney U-test (**D-E)** Representative MIP images of adult fly antennal lobe expressing CaMPARI and Ir64a RNAi in nSyb-expressing neurons exposed to 5% AA at pH=2 **(D)** or pH=7 **(E)** for 30 min. Photoconversion is not seen in the red channel when the pH=2 **(D)** or pH=7 **(E). (F)** The Red:Green ratio of flies exposed to AA of pH=2 and pH=7 for 30 minutes. Flies exposed to AA pH=2 (n=14) have 1.3-fold greater photoconversion compared to flies exposed to AA pH=7 (n=15). *P=0.0151, Mann-Whitney U-test

To further confirm the specificity of photoconversion in flies expressing pan-neuronal CaMPARI, these flies were crossed with a line capable of expressing a shRNA directed at Ir64a mRNA, thus triggering RNA interference (RNAi), to induce knockdown of the acid-sensing Ir64a receptor. Three sets of 4-5 flies were exposed to 5% acetic acid or neutralized 5% acetic acid for 30 minutes while simultaneously exposed to photoconvertible light (405 nm; 500 ms on, 200 ms off). The brains were then dissected and imaged as described above. Ratios were calculated from AIP images as described above. We found that freely-moving intact adult flies with pan-neuronally expressing CaMPARI and Ir64a RNAi exposed to 5% AA (pH=2) (n=14 individual flies) (**Figure 3D)** had 1.3-fold photoconversion compared to flies exposed to 5% AA (pH=7) (n=15 individual flies) (**Figure 3E**) in the DC4 glomerulus (**Figure 3F**). Even with knockdown of Ir64a, the amount of photoconversion in flies exposed to 5% AA (pH=2) is still significantly increased compared to flies exposed to neutralized AA, however the amount of photoconversion is reduced by 50% compared to flies not expressing the Ir64a RNAi. This is probably due to incomplete knockdown of the receptor. Still, this reduction in photoconversion suggests the photoconversion seen when flies are exposed to 5% AA (pH=2) is specific to activation of neurons expressing the Ir64a receptor.

### 4.4 Apple cider vinegar exposure results in specific photoconversion of the DC4 glomerulus in flies expressing CaMPARI

We next wanted to compare the photoconversion we demonstrated with 5% acetic acid to a similar physiologically-relevant stimuli, apple cider vinegar, which is typically attractive to *Drosophila melaogaster*. Five to six sets of 3-5 flies expressing CaMPARI driven by a synaptobrevin (nSyb) GAL4 driver or an Ir64a GAL4 driver were exposed to apple cider vinegar (ACV) or neutralized ACV for 30 minutes while simultaneously exposed to photoconvertible light (405 nm; 500 ms on, 200 ms off) and their brains dissected and imaged as described above. Maximum intensity projection (MIP) images of one antennal lobe are shown in **Figure 4** for flies exposed to either 5% AA (pH=2) (**Figure 4A** and **4C**) or neutralized 5% AA (pH=7) (**Figure 4B** and **4D**). Ratios were calculated from AIP images as described above. We found that freely-moving intact adult flies with pan-neuronally expressing CaMPARI exposed to ACV (pH=2) (n=27 individual flies) had 2.1-fold photoconversion compared to flies exposed to neutralized ACV (pH=7) (n=21 individual flies) in the DC4 glomerulus (**Figure 4E**). Similarly, flies expressing CaMPARI specifically in Ir64a-expressing neurons exposed to ACV (pH=2) (n=19 individual flies) had 3.1-fold photoconversion compared to flies exposed to neutralized ACV (pH=7) (n=18 individual flies) (**Figure 4F**).

**Figure 4:**
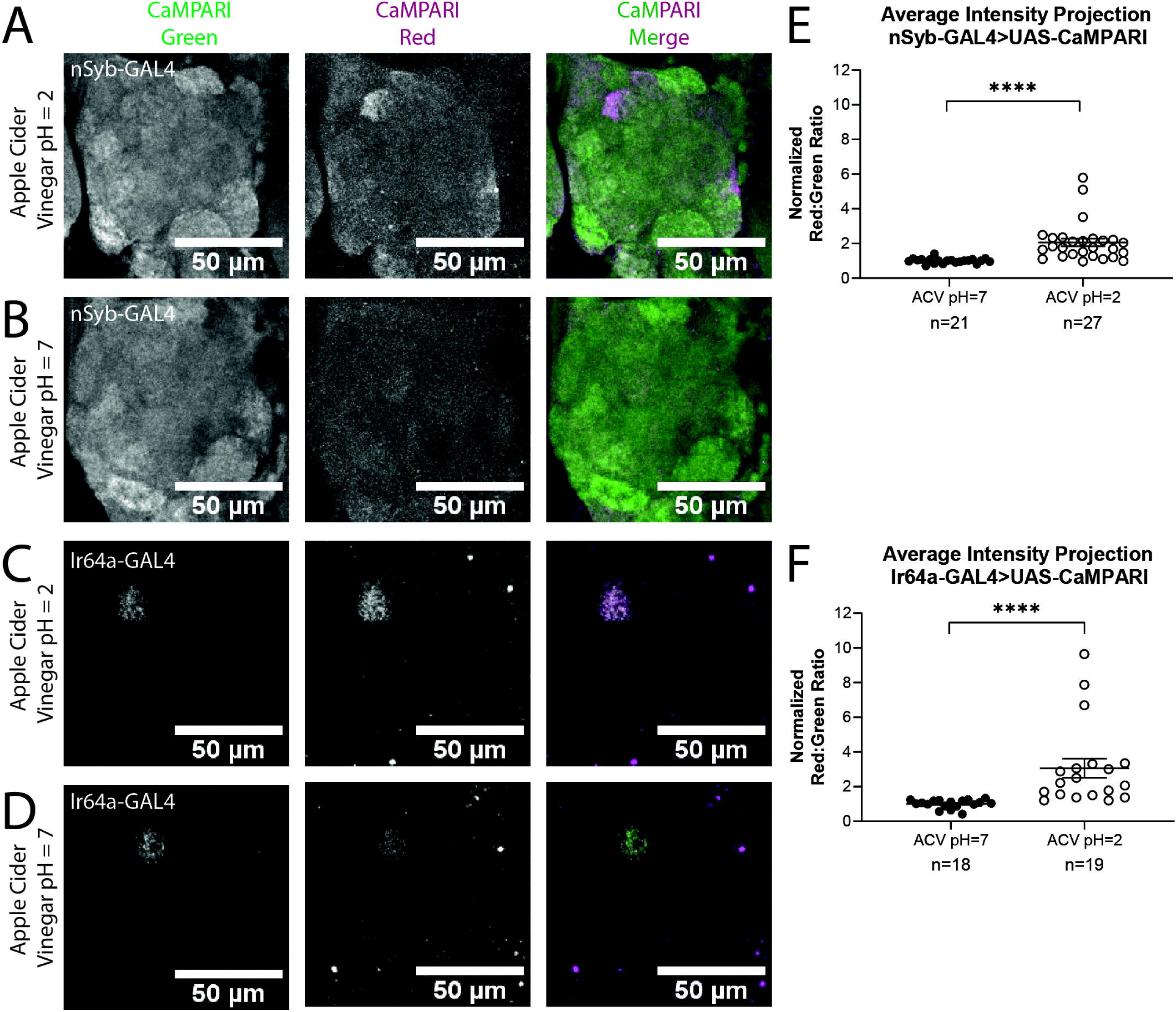
Flies Expressing CaMPARI Have Measurable Photoconversion When Exposed to Apple Cider Vinegar. **(A-B)** Representative maximum intensity projection (MIP) images of adult fly antennal lobe expressing CaMPARI in nSyb-expressing neurons exposed to 5% apple cider vinegar (ACV) at pH=2 **(A)** or pH=7 **(B)** for 30 minutes. Photoconversion is present in Ir64a-expressing neurons in **(A)** when the pH=2 however no appreciable photoconversion is seen in **(B)** when the pH=7. **(C-D)** Representative MIP images of adult fly antennal lobe expressing CaMPARI in Ir64a-expressing neurons exposed to 5% ACV at pH=2 **(C)** or pH=7 **(D)** for 30 minutes. Photoconversion is seen in the red channel in **(C)** when pH=2 but not in **(D)** when pH=7. **(E)** The Red:Green ratio of flies exposed to ACV of pH=2 and pH=7 for 30 minutes. Flies exposed to ACV pH=2 (n=27) have 2.1-fold greater photoconversion compared to flies exposed to ACV pH=7 (n=21). ****P<0.0001, Mann-Whitney U-test **(F)** The Red:Green ratio of flies exposed to ACV of pH=2 and pH=7 for 30 minutes. Flies exposed to ACV pH=2 (n=27) have 3.1-fold greater photoconversion compared to flies exposed to ACV pH=7 (n=21). ****P<0.0001, Mann-Whitney U-test

## 5 Discussion

Previously, researchers have used intermediate early genes (IEGs), such as Arc (Link et al., 1995; Guzowski et al., 1999) and cFos (Cole et al., 1989) as an indirect measure of neuronal activation, however the use of IEGs is limited by the transient nature of their expression and lack of temporal resolution (Fields et al., 1997). An additional commonly used tool is genetically encoded calcium indicators (GECIs), such as GCaMP (Riemensperger et al., 2012). GCaMP is a powerful tool for visualizing immediate activation but usually limits the movement of the organism during imaging and also limits the field of view during imaging. CaMPARI was developed as a solution to these limitations, however methods requiring the adult fly to be fixed underneath a scope have continued to limit its usefulness in studying the complex behaviors of adult *Drosophila*.

We have developed a method to trace active neural circuitry in freely-moving intact adult *Drosophila*. While previous methods have used direct laser light from microscopes, requiring immobilization of the fly (Bohra et al., 2018; Manjila et al., 2019), our method using LEDs and Petri dishes allows the flies to remain mobile and freely behaving and allows us to capture the active circuitry of multiple flies simultaneously. We demonstrate that this method faithfully recapitulates a previous study (Ai et al., 2010) demonstrating that exposure to acids activates the Ir64a-expressing neurons of the DC4 glomerulus in the antennal lobe of adult flies.

Using CaMPARI specifically expressed in Ir64a-expressing neurons while exposing flies to acetic acid, we showed that we can label active neurons in freely-moving adult flies and quantify the photoconversion in the DC4 glomerulus. The red:green ratio of flies exposed to acetic acid was significantly higher than flies exposed to neutralized acetic acid. Additionally, we showed that photoconversion between the left and right sides of the brain was fairly consistent, suggesting that light penetration from the LED was evenly distributed to both sides of the head. We were able to recapitulate these results with a pan-neuronally expressed CaMPARI. Quantification of the photoconversion in these flies was equivalent to the photoconversion seen when CaMPARI expression was limited to Ir64a-expressing neurons. This indicates CaMPARI can be used quantitatively whether CaMPARi expression is limited to subsets of neurons or is expressed broadly. CaMPARI expression is only limited by the strength and specificity of available driver lines. The ability to faithfully label active neurons with pan-neuronally expressing CaMPARI suggests CaMPARI can be used to label active circuitry even without *a priori* knowledge of which neurons should be active with a stimulus. Thus, our method allows for CaMPARI to be used as an exploratory tool in freely moving flies to discover novel parts of a circuit.

While our results show promise for CaMPARI as an exploratory tool in freely moving flies, there remain some caveats which may be addressed as the tool continues to develop. For example, the signal of red CaMPARI can be relatively low and in some low-activity circuits may be indistinguishable from background. Two improvements to CaMPARI have already been made to address these concerns: (1) an improved version of CaMPARI (CaMPARI2) that improves the brightness and contrast of red CaMPARI and (2) an antibody that specifically detects the red CaMPARI chromophore, but not the green chromophore (Moeyaert et al., 2018). With these tools, CaMPARI freely diffuses through the cytoplasm and so subcellular information is lost. Recently the tool SynTagMa (Synaptic Tag for Mapping Activity) was created to label active synapses by fusing CaMPARI to synaptophysin or PSD95 and labeling presynaptic boutons and dendritic spines, respectively (Perez-Alvarez et al., 2019). This tool could be further adapted to more specifically label circuitry in adult *Drosophila* or any other species where transgenic methods allow creation of viable transgenic animals. Additionally, pairing CaMPARI with other optogenetic tools will allow downstream tracing of active circuits. For example, using CsChrimson to activate a specific neuron while exposing a pan-neuronal expressed CaMPARI will allow an experimenter to trace an neurons excited by the neuron expressing CsChrimson. Developing a method that allows whole-brain tracing of active circuitry in freely moving adult flies can allow CaMPARI to be used in an unbiased and exploratory manner. It allows for study of various behaviors from movement to sensory response to social interaction and beyond.

## 6 Conflicts of Interest

The authors declare that the research was conducted in the absence of any commercial or financial relationships that could be construed as a potential conflict of interest.

## 7 Author Contributions

KE, MB and GB designed the study. KE performed the experiments and analyzed the data. KE, MB and GB wrote the paper. All authors read and edited the manuscript.

## 8 Funding

This project was supported by grants from the Geisel School of Medicine at Dartmouth (GB), the National Institute of Health Pioneer grant 1DP1MH110234 (GB), the Defense Advanced Research Projects Agency grant HR0011-15-1-0002 (GB), and the National Institute of General Medical Sciences grant GM113132 (MBH).

## 9 Acknowledgments

We thank the Bloomington Drosophila Stock Center and Vienna Drosophila Resource Center for stocks. We thank Vibhuti Rana for careful reading of the manuscript and Balint Kacsoh and Madhu Sadanandappa for helpful comments as the project progressed. We would also like to thank Ann Lavanway and the Dartmouth College Department of Biological Sciences Imaging Facility for the use of equipment and technical assistance as well as Zdenek Svindrych of the Molecular Interactions and Imaging Core for technical assistance.

