## Supplemental Figures for "The Photoconvertible Fluorescent Probe, CaMPARI, Labels Active Neurons in Freely-Moving Intact Adult Fruit Flies"

***Supplementary Material***

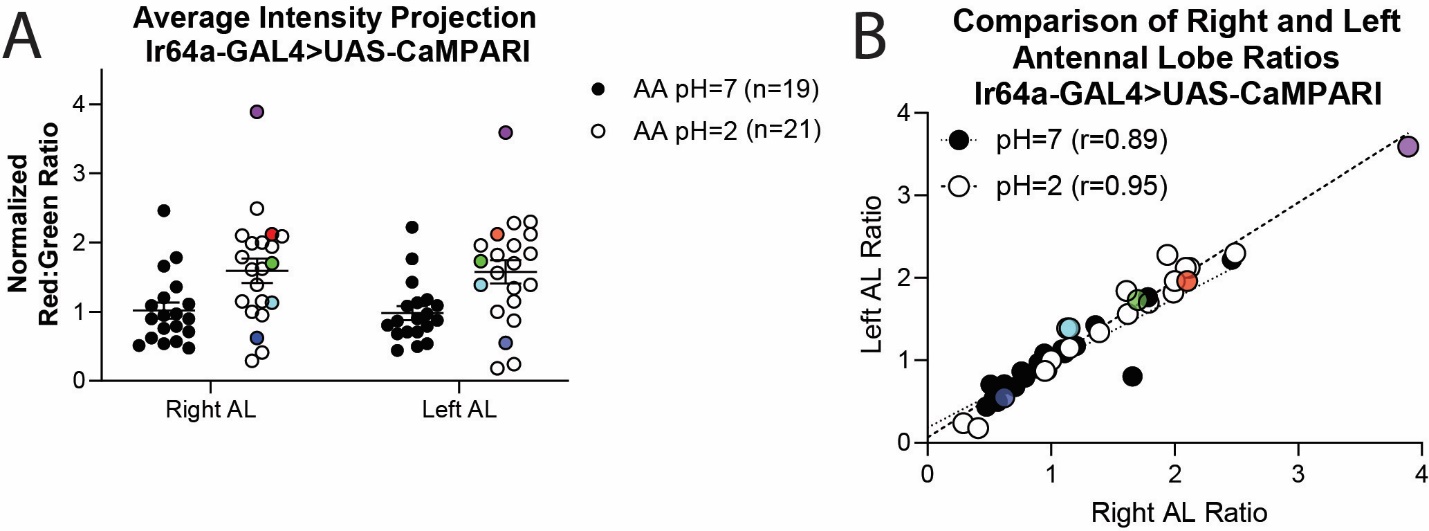
**1	Supplementary Figures**

**Figure S1: CaMPARI Photoconversion Is Comparable Between Right and Left Glomeruli Measured from the Same Brain**

**(A)** The Red:Green ratio of flies exposed to AA of pH=2 and pH=7 for 30 minutes. Individual measurements are shown for each right and left antennal lobe (AL) of an adult fly brain average intensity projection. Five color-matched paired points demonstrate that measurements taken from both glomeruli of the same brain are similar and so have been averaged together in subsequent graphs to give one point for each fly brain.**(B)** A direct comparison of the Red:Green ratios for each right and left AL demonstrates that measurements taken from the same brain are comparable. Right and left AL are correlated with a slope of 0.78 and r=0.89 for flies exposed to neutralized acetic acid and a slope of 0.95 and r=0.95 for flies exposed to acetic acid (pH=2).

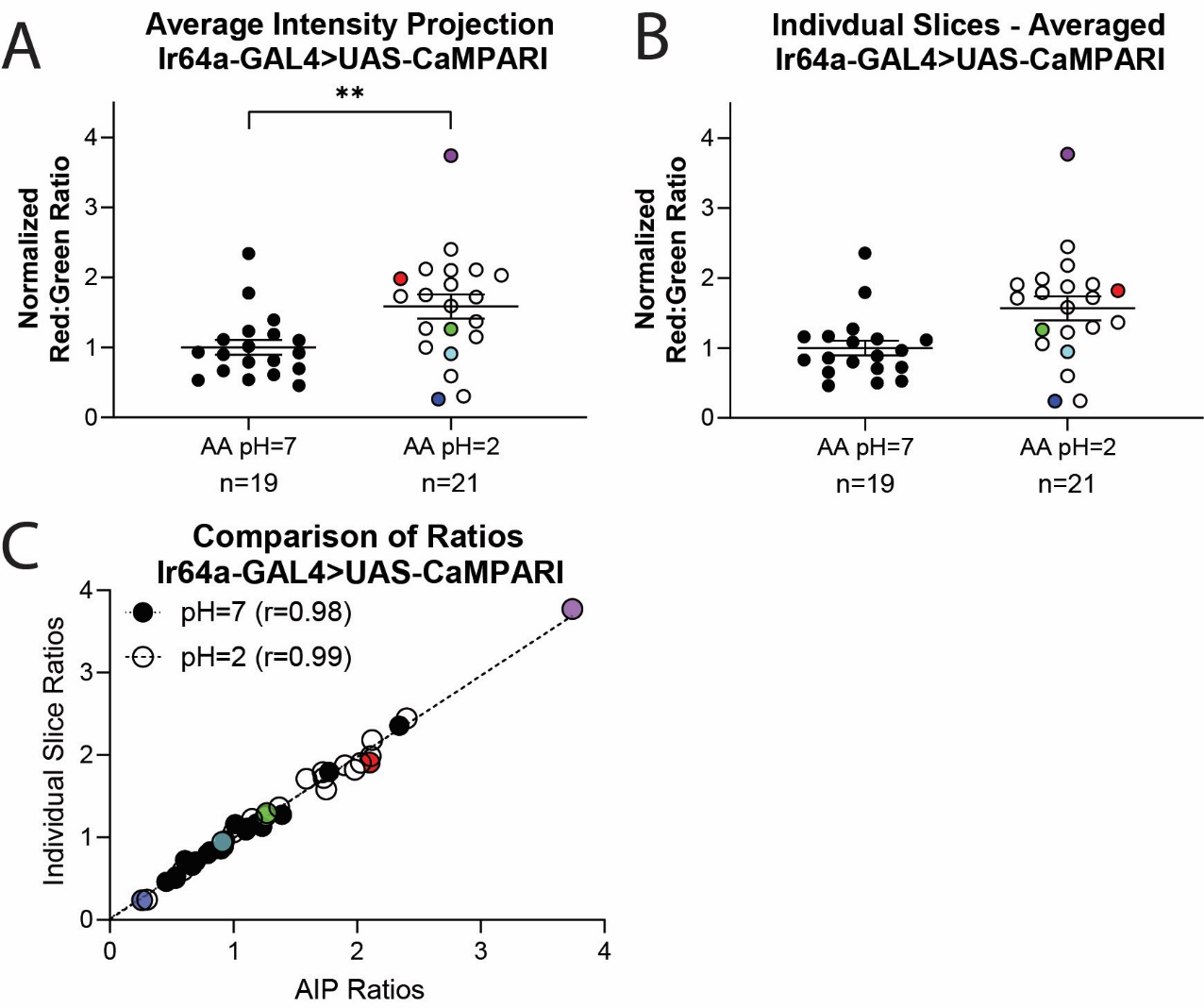

**Figure S2: Photoconversion Ratios Measured from Average Intensity Projection Images are Similar to Measurements from Individual Slices Averaged Together**

**(A-B)** The Red:Green ratio of flies exposed to AA of pH=2 and pH=7 for 30 minutes from adult fly brain average intensity projection (AIP) images **(A)** or individual slices containing the region of interest averaged together **(B)**. Five color-matched paired points demonstrate that measurements taken from AIP images are similar to measurements taken from images of the same fly brain where individual slice measurements have been averaged together. All subsequent graphs show measurements taken from AIP images. **(C)** A direct comparison of the Red:Green ratios for AIP measurements and individual slice measurements demonstrating measurements taken from the same brain are comparable. Right and left AL are correlated with a slope of 0.98 and r=0.98 for flies exposed to neutralized acetic acid and a slope of 0.98 and r=0.99 for flies exposed to acetic acid (pH=2).
